# DNAJB6b — a complete amyloid inhibitor

**DOI:** 10.64898/2026.09.03.749188

**Authors:** Andreas Carlsson, Ulf Olsson, Sara Linse

## Abstract

In Alzheimer’s disease (AD), the aggregation of amyloid β (Aβ) peptides has been linked to disease pathology. Once aggregates have formed, existing fibrils catalyze the formation of new fibrils in a runaway process associated with neuro degeneration. One component of the cellular defense system against amyloid formation is the molecular chaperone DNAJB6b (JB6). JB6 is known to suppress both primary nucleation and fibril-catalyzed secondary nucleation. However, the extent of JB6 binding to Aβ fibrils and whether this interaction leads to inhibition of fibril elongation also, have not been established. Here, we combine well-characterized aggregation kinetics of Aβ42 and the knowledge of JB6 self-assembly equilibrium and kinetics, with high-sensitivity HPLC measurements of free JB6 at nanomolar concentrations to determine the affinity of JB6 for Aβ42 fibrils. We show that JB6 binds to Aβ42 fibrils with high affinity and that this binding can quantitatively explain the observed elongation inhibition. Diffusion-based techniques reveal that JB6 binds to amyloid intermediates, resulting in potent suppression of amyloid nucleation until the pool of free JB6 is depleted, after which rapid bulk aggregation ensues. Our results provide a mechanistic link between chaperone–fibril binding and amyloid suppression, offering a quantitative framework for understanding how molecular chaperones regulate amyloid formation.

## Introduction

DNAJB6b (JB6) is one of the most effective chaperones in amyloid suppression [1, 2, 3]. JB6 has been found to suppress amyloid formation of several peptides both *in vivo* and *in vitro*, including α-synuclein [4, 5, 6, 7, 8], polyglutamine peptides [1, 9, 10, 11, 12, 13], tau [2, 3, 14], amyloid β (Aβ) [11, 15, 16, 17, 18, 19], TDP43 [20], FUS [21], and phenylalanine-glycine-rich nucleoporins [22]. In Alzheimer’s disease (AD), the aggregation of Aβ peptides precedes disease-related events such as tau aggregation, which leads to neurodegeneration, and cognitive decline [23]. JB6 is predominantly found intracellularly [1, 7, 24], whereas Aβ is more commonly found extracellularly. However, there is a growing body of findings of Aβ entering cells as well [25, 26, 27], highlighting the importance of characterizing the suppression of Aβ aggregation by JB6. Whether JB6 also enters the extracellular space remains unknown, although the chaperone has been identified in extracellular vesicles [28]. The evolutionary emergence of homologs to JB6, or other JDPs of the non-canonical B class, has been found to precede the emergence of amyloid prone peptides, further suggesting an important role in suppressing amyloid formation [29].

Aβ42 is the dominating peptide in plaques in AD [30], and this peptide is the focus of the present work. The aggregation process of Aβ (especially by the 40- and 42-residue peptides) has been extensively studied, revealing that secondary nucleation is the dominant process for the generation of new fibrils [31, 32, 33], whereas elongation is the process generating most fibril mass and primary nucleation is required for initial aggregation from monomers. The rate constants for the microscopic steps in Aβ40 and Aβ42 aggregation have been estimated [34].

With JB6 we refer to isoform b of the human DNAJB6. Isoform a is found in the nucleus [24] and will not be considered in the present work. JB6 has an N-terminal J-domain, which can bind to HSP70 and initiate ATP-dependent pathways leading to a decrease in protein aggregates, via refolding [35], proteasomal degradation [36], or fragmentation followed by lysosomal degradation [37]. A first, critical step in this disease-preventive network is for JB6 to bind to the aberrant protein aggregates, analogous to an antibody recognizing a specific pathogen. This pivotal interaction between JB6 and protein aggregates seems to be independent of other cellular components and is therefore well suited for *in vitro* studies, where components, concentrations, and solution conditions can be precisely controlled.

JB6 exists in a self-assembly equilibrium between monomers [38] and micellar-like oligomers, with a critical micelle concentration (cmc) of approximately 120 nM [39]. Importantly, the monomeric form is substantially more potent in suppressing amyloid formation than the micellar assemblies [18]. The dissociation is strongly temperature dependent, with a half-time of micelle dissociation of approximately 20 min at 37 °C [38]. Thus, the chaperone activity of JB6 depends not only on its concentration but also on its assembly state. This becomes particularly important when studying rapid amyloid processes, such as fibril elongation in highly seeded reactions, where aggregation may occur on a timescale comparable to or faster than JB6 micelle dissociation [18]. Accounting for this self-assembly behavior therefore allows us to isolate the activity of monomeric JB6 and determine its effect on fibril elongation.

In the present work, we investigate the extent to which JB6 inhibits Aβ42 fibril elongation. JB6 has previously been shown to suppress both primary and fibril-catalyzed secondary nucleation of Aβ42 [11]. Here, we mainly focus on the elongation, the microscopic process that generates fibril mass. We further quantify the affinity and stoichiometry of JB6 binding to Aβ42 fibrils and ask whether these binding parameters can quantitatively explain the inhibition of fibril growth in seeded reactions. Finally, we investigate the interaction of JB6 with transient amyloid species during aggregation to learn how chaperone availability evolves throughout the reaction. Together, these experiments provide a quantitative framework linking the molecular association of JB6 with Aβ42 species to its inhibition of the major microscopic processes of amyloid formation.

## Results and Discussion

### Inhibition of seeded Aβ42 aggregation by JB6

JB6 has previously been shown to inhibit both primary and secondary nucleation of Aβ42 [11], but whether elongation — the process generating most of the fibril mass — is also affected has remained unresolved. One reason for this has been the lack of background knowledge required to design conclusive experiments. It was only recently established that JB6 needs to dissociate into monomers in order to become fully active [18], a process that occurs on a time scale of approximately 20 minutes at 37 °C [38], which is slow relative to the rapid aggregation in highly seeded samples typically used when examining elongation. Thus, when JB6 is diluted from a concentrated stock to low nanomolar concentrations immediately before the reaction, the micelles do not have sufficient time to dissociate before most of the aggregation process is over.

To address the effect on elongation, JB6 was therefore here first equilibrated at near its final concentration, after which seeds were added to allow the chaperone to bind, followed by addition of monomeric Aβ42. Aggregation was probed using two different techniques, thioflavin T (ThT) fluorescence and circular dichroism (CD) spectroscopy. Measurement of ThT fluorescence with a plate reader enables high-throughput measurements, while CD spectroscopy probes amyloid formation without any added fluorophore, providing a direct measurement of the β-sheet content.

Primary and secondary nucleation was here bypassed by starting the aggregation reaction with 5 % sonicated seeds and a total Aβ42 concentration of 5 µM. Under this condition, the nucleation of additional fibrils can to a good approximation be neglected. Having elongation only, the kinetics is described by a single exponential function (*f* (*t*) = 1 *−* 0.95*e^−kt^*) [40] in the case of normalized data with 5 % seeds. Figure 1a shows a single exponential fit to the data of Aβ42 alone, with the rate constant *k* as the only free parameter. As can be seen, the single exponential function describes the data well, and from the fit we obtain *k* = 0.00182*±*0.00009 s*^−^*^1^.

**Figure 1:**
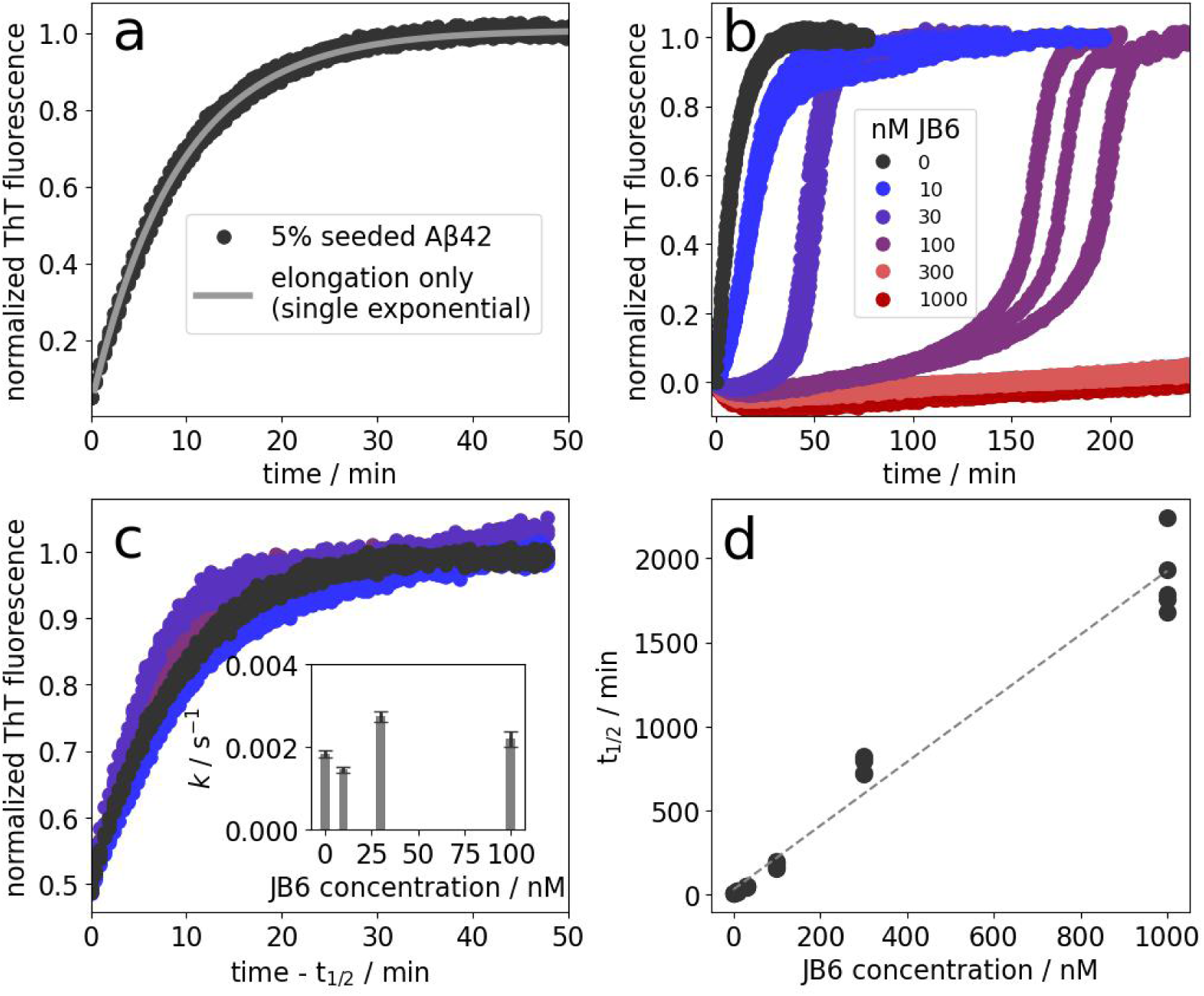
The effect of JB6 on Aβ42 elongation, probed with ThT fluorescence. **a**: Aggregation kinetics of Aβ42 alone (5 % sonicated seeds, 5 µM total Aβ42) with a fit assuming growth via elongation only: f (t) = 1 − 0.95e−kt. b: The effect of JB6 on elongation was studied using 0.25 µM (5 %) sonicated Aβ42 seeds, with a total Aβ42 concentration of 5 µM and five replicates at each JB6 concentration. c: Comparison of the final 50 % of the kinetic traces. The inset shows the mean values and standard deviations (n = 5) for fits of k using f (t) = 1 − 0.5e−kt. d: t1/2-values from traces in (a) and (b), plotted versus the JB6 concentration. The dashed line is a linear fit to the data points.

*k* is related to the elongation rate constant, *k*_+_, by *k* = *k*_+_2[*P*], where [*P*] is the number concentration of fibrils, and 2*P* is the number concentration of fibril ends. With reasonable assumptions of the number of monomers constituting a fibril [41, 42, 43, 44], the *k*_+_ is found to be similar to previously findings [34] (see supplemental note 1 for the calculations).

The aggregation of Aβ42 is strongly suppressed by JB6 in a manner dependent on the JB6 concentration (Figure 1b). Since elongation determines the aggregation kinetics under these seeding conditions, it is evident that JB6 diminishes the elongation rate of Aβ42. The initial slope approaches zero at higher JB6 concentration, consistent with near-complete inhibition of the elongation process. However, the second half of the aggregation reaction proceeds with similar kinetics independent of the JB6 concentration, as shown in panel (c), plotted by shifting the time axis to start at the half-time. This is in line with what has earlier been observed [11]. The traces were fitted to an elongation-only model in the supplementary information, SI, (Figure S1) and the obtained *k*-values are plotted in the inset of panel (c). It may be noted that Aβ42 eventually aggregates even in the presence of 300 and 1000 nM JB6 (Figure S2a) but we focus here on the JB6 concentrations below its cmc to avoid complications arising from JB6 self-assembly and the possibility that both JB6 monomers and micelles bind to the fibrils. Furthermore, the retardation increases approximately linearly with the JB6 concentration, as seen in Figure 1d, where the halftime of aggregation, t_1_*_/_*_2_, is plotted against the JB6 concentration.

ThT fluorescence is often used as a reporter of fibril mass, but it is well known that the quantum yield of the bound dye depends strongly on the fibril structure and environmental conditions, which may result in misleading kinetic traces [45, 46]. In addition, the ThT may compete with JB6 in fibril binding, and ThT displays a weak fluorescence in the presence of JB6 alone (Figure S2b), which may further complicate the interpretation as the extent of chaperone-binding may shift the signal. To probe the aggregation with a more direct measure, circular dichroism (CD) spectroscopy was used, monitoring the change in ellipticity associated with β-sheet formation. This way of probing the amyloid formation is especially advantageous when analyzing the first minutes of the kinetic trace where temperature variations may skew the ThT fluorescence signal. While the CD signal will report also on β-sheet in the chaperone, this is likely a constant contribution over time and in all cases the chaperone concentration is much lower than the Aβ42 concentration.

Figure 2a shows the CD spectrum variation during an aggregation reaction starting from 12 µM monomeric Aβ42. The ellipticity at 218 nm is used as a reporter of the aggregation process and the value of this parameter becomes more negative over time reflecting increasing fibril mass concentration. A typical sigmoidal curve shape is obtained, as is expected for a nucleated process, and the relatively steep transition reveals the existence of a secondary nucleation step in addition to primary nucleation and elongation [47].

**Figure 2:**
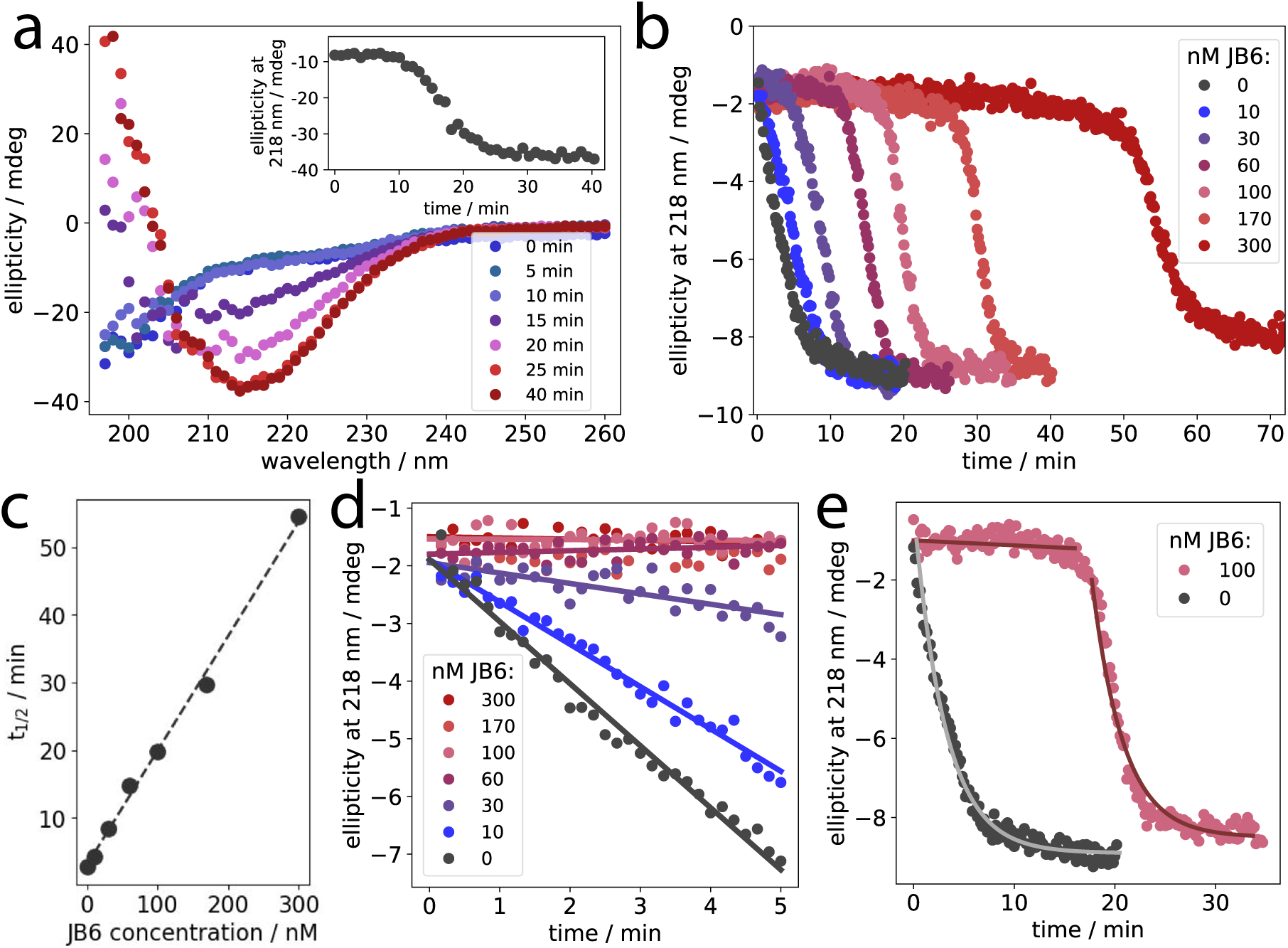
CD spectroscopy to probe Aβ42 aggregation kinetics. **a:** Time-resolved CD spectra of the conversion from unfolded, monomeric Aβ42 (12 µM), into β-sheet-rich amyloid. Inset: Ellipticity at 218 nm during the fibril formation process. b: β-sheet formation through ellipticity at 218 nm, with a starting point of 5 % sonicated seeds, a total Aβ42 concentration of 5 µM, and varying JB6 concentrations. c: The half time, t1/2, versus JB6 concentration, with a linear fit shown as a dashed line. d: First 5 minutes of the kinetic traces from panel (b). Linear regressions are shown as solid lines. e: Describing the kinetic trace of Aβ42 alone with a single exponential decay function, with a k-value referred to as k0, and the trace corresponding to the presence of 100 nM JB6 with two phases: the first with k=0.005k0 and the second with k0.

The addition of 0.25 µM sonicated seeds to 4.75 µM monomeric Aβ42 (i.e. 5 % seeds) makes both primary and secondary nucleation bypassed, since elongation becomes the dominant microscopic process and proceeds rapidly, as shown in Figure 1a. Under these conditions, the fibril mass concentration initially increases linearly with time and then gradually approaches a plateau, i.e., the inverse of the exponential decay of free monomer, as observed for Aβ42 alone in Figure 2b (black dots). Upon addition of JB6 at the start of the reaction, the initial slope decreases with increasing JB6 concentration and becomes nearly flat at sufficiently high JB6 concentrations, consistent with a strong inhibition of elongation. CD spectra on the end-states are highly similar (Figure S3).

Similar to the aggregation kinetics monitored by ThT fluorescence in a PEGylated polystyrene plate, JB6 delays the aggregation approximately linearly with JB6 concentration, as seen in the t_1*/*2_, Figure 2c.

It may be noted that although the kinetic traces obtained with ThT fluorescence and CD spectroscopy have similar shapes, the absolute timescales differ considerably. This is not surprising since both sample mixing during the reaction and container surfaces differ greatly between the two cases [43]. The ellipticity were measured in a quartz cuvette with continuous stirring (magnetic stir bar) to sustain a homogeneous dispersion, whereas the samples in PEGylated 96-well plates were monitored without shaking, in which case the only agitation imposed on the samples comes from the continuous moving of the plate over the plate reader optics.

Since the aggregation is dominated by elongation with 5 % sonicated seeds, the initial part of the trace is linear and the slope corresponds to the elongation rate. The first 5 minutes of the aggregation traces in Figure 2b are analyzed with linear regressions, shown in panel (d) to obtain how the elongation rate depends on JB6 concentration. Using a function that only considers elongation (single exponential decay), the *k*-value may be varied during the aggregation to describe the aggregation kinetics. For Aβ42 alone, only one *k*-value, called *k*_0_, should be needed to describe the whole reaction, which is the case as seen in Figure 2e. In the presence of 100 nM JB6, the lag phase has a slope that is determined in (d) to be 0.005*k*_0_. The two red lines in (e) corresponds to elongation growth with first *k* = 0.005*k*_0_ and then *k*_0_. Thus, after the apparent lag phase, the aggregation kinetic is highly similar to that of Aβ42 alone, in agreement with the ThT fluorescence measurements (Figure 1c).

The aggregation kinetics experiments using both ThT fluorescence and CD spectroscopy outline a consistent picture: JB6 inhibits Aβ42 elongation, resulting in lag phases with increasing length with increasing JB6 concentration. After the lag phase, the aggregation of Aβ42 proceeds with kinetics similar to that observed in the absence of JB6. Månsson et al. [11] suggested that this scenario reflects JB6 being “consumed” by associating to the growing fibrils. To understand this behavior quantitatively, we measured the binding isotherm of JB6 to Aβ42 fibrils as presented below.

### The affinity of JB6 for Aβ42 fibrils

The affinity of JB6 for Aβ42 fibrils is fundamental to our understanding of chaperone action by JB6. To measure the fibril-binding affinity, we utilized the high sensitivity of HPLC (reversed phase with UV-absorbance) to quantify JB6 concentrations down to below 1 nM. The studies of binding events involving monomeric JB6 was enabled by working in a concentration range well below the cmc of JB6 (which is about 120 nM [39]) and by giving JB6 sufficient time to dissociate into monomers [18, 38] before the addition of fibrils. Figure 3 shows binding data obtained by incubating monomeric JB6 with preformed Aβ42 fibrils and measuring the free JB6 concentration after a centrifugation step to remove fibrils with bound JB6 (see Figure S4a for an investigation of centrifugation times).

**Figure 3:**
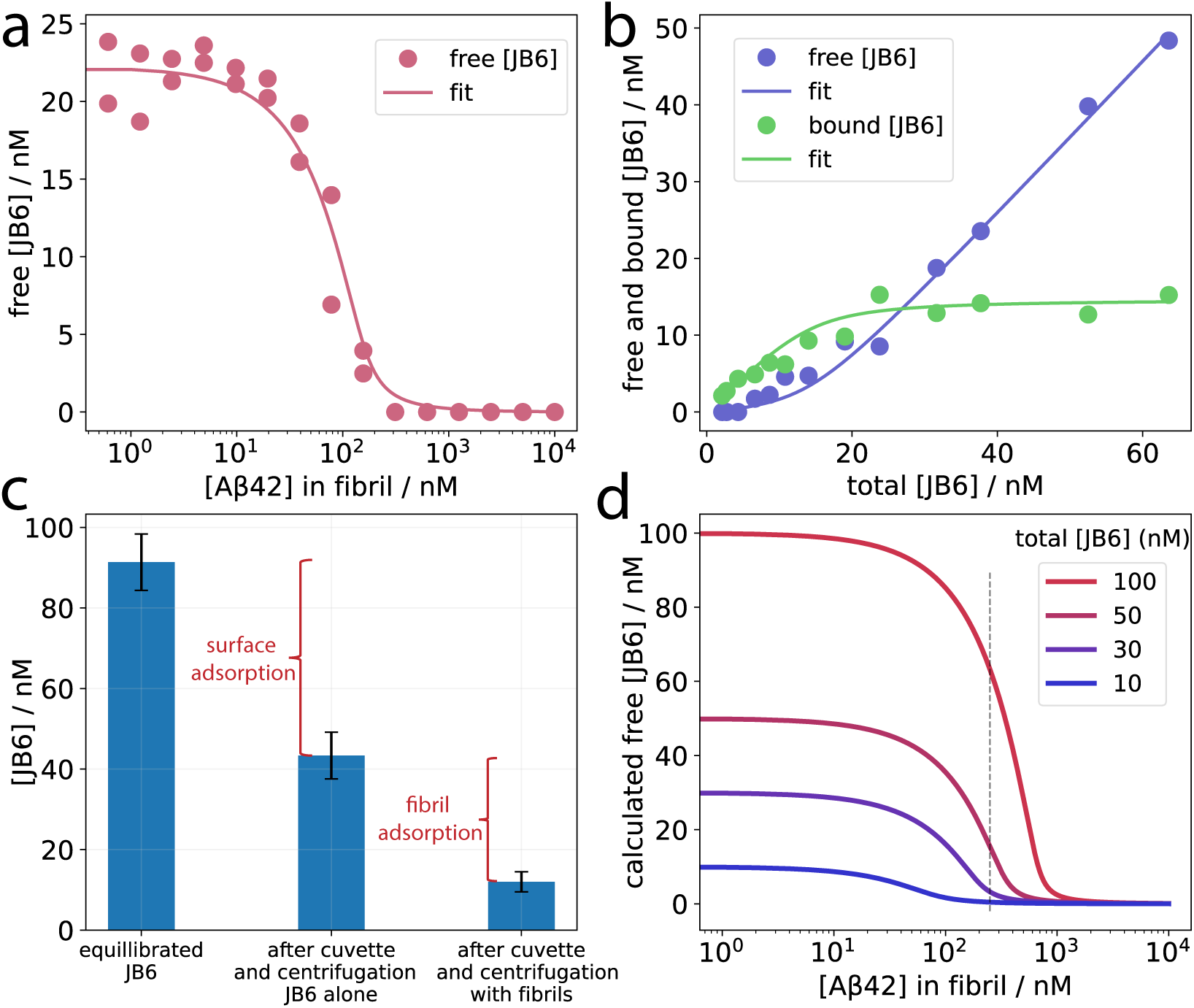
Estimation of the affinity of JB6 for Aβ42 fibrils. **a:** The concentration unbound (free) JB6 was measured in samples with constant JB6 concentration (22 nM) and varying Aβ42 fibril concentration. b: Varying JB6 concentration and keeping the fibril concentration constant (100 nM in monomer equivalents). The free JB6 concentration is measured and the bound JB6 concentration is calculated as total minus free. A global fit to all data in panel (a) and (b) yields a KD of about 1 nM and a saturation stoichiometry of approximately 1:7 JB6 to Aβ42 monomers in the fibril state. c: For the estimation of how much JB6 is bound to fibrils, surface adsorption of JB6 to the container needs to be accounted for. This is showcased here with an example of 100 nM JB6 and 250 nM Aβ42 fibrils. An identical sample with only JB6 is used to measure how much JB6 is bound to container surfaces, before the addition of fibrils. The error bars represent standard deviation of n=3 measurements. Methodological details are found in SI, supplemental note 2. d: Calculated concentrations of free JB6 versus fibril concentration, using the affinity parameters obtained in (a) and (b). The dashed line at 250 nM fibril corresponds to the seed concentration used in the aggregation kinetics experiments.

In the experiment shown in Figure 3a, the JB6 concentration is kept constant while the fibril concentration is varied. Note that the fibril concentration is defined in monomer equivalents. The data are well described by a binding isotherm for independent binding, suggesting a lack of cooperativity. The data moreover display a decent coverage of the relevant concentration range, providing high sensitivity for the affinity of the interaction.

In another experiment, shown in Figure 3b, the JB6 concentration is varied while the fibril concentration is kept constant, generating a complementary data set sensitive to the maximum amount of bound JB6 and thereby allowing determination of the binding stoichiometry. Working with surface active proteins at low concentrations requires careful consideration of container material and sample volumes. JB6 is highly surface active [14, 48], and to accurately obtain the JB6 amount bound to fibrils, it is important to measure the available concentration of JB6 in the samples (used in Figure 3b on the x-axis), which was here done in identical samples without fibrils. This concentration is the accessible JB6 concentration since JB6 was equilibrated in the tubes before the addition of fibrils and JB6 surface adsorption is practically irreversible over the time-scale of the experiment [48]. Panel (c) showcases an example of how much JB6 is adsorbed at container surfaces and fibrils in the case of 100 nM JB6 added to 250 nM fibrils. Supplemental note 2 shows calculations to relate the adsorbed amount to the affinity measurements.

Binding parameters were obtained from a global fit of a binding isotherm to both data sets:

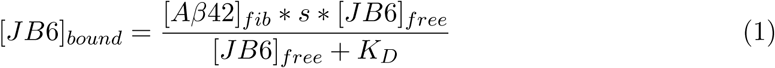

with

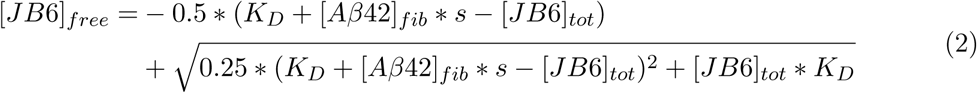

where [*JB*6]*_free_* is the free JB6 concentration, [*Aβ*42]*_fib_* is the fibril concentration in monomer equivalents, and [*JB*6]*_tot_* is the total JB6 concentration. *K_D_* is the equilibrium dissociation constant, fitted to 1.3 nM (0-2.6 nM 95% confidence interval), and *s* is a stoichiometry factor at saturation, fitted to 0.15 (0.13-0.16 95% confidence interval), corresponding to approximately one JB6 molecule per seven Aβ42 molecules. The affinity is high and similar to the affinity of JB6 for amyloid oligomers of an Aβ fragment (20-34) [19]. The affinity and stoichiometry fairly similar to that measured for fibrils formed of an amyloid core fragment of tau (*K_D_* = 4.7 nM, s = 0.1) [14]. The methodology may be used to reveal whether JB6 is specific for amyloid fibrils, or binds to other types of solid aggregates as well.

Using the experimentally obtained affinity and stoichiometry, the concentration of free JB6 can be calculated for any given fibril concentration. This is illustrated with a few example concentrations in Figure 2d.

### Linking chaperone-fibril binding to the inhibition of amyloid formation

The potency of the amyloid inhibition may be interpreted with the fibril-affinity of monomeric JB6. We do this by comparing the elongation inhibition with the JB6-coverage of the fibrils at time zero in the aggregation reaction (Figure 4).

**Figure 4:**
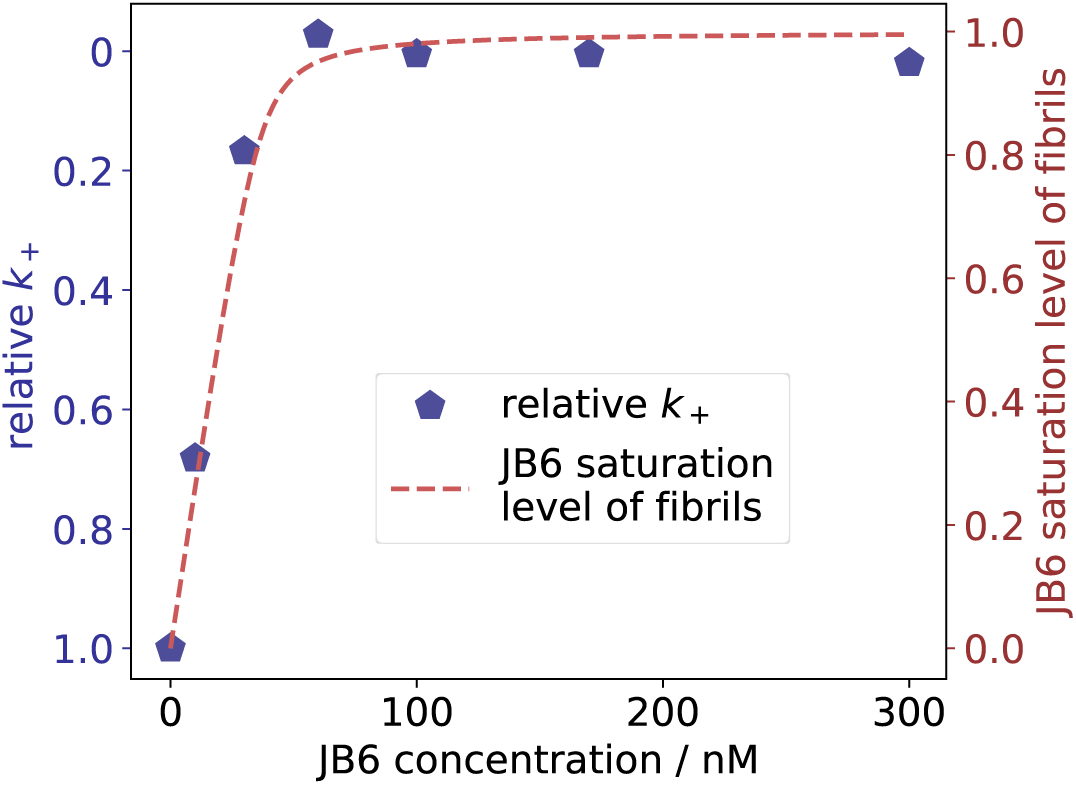
Coupling amyloid inhibition with fibril-affinity. The initial slopes in Figure 2d report on the elongation rates. By dividing each slope by the slope for Aβ42 alone, the relative elongation rate constants, k+, are obtained, shown as blue pentagons on the left y-axis. The fibril coverage by JB6, called “JB6 saturation level of fibrils” is calculated for the initial state of the aggregation reactions with 250 nM fibrils (seeds), for each JB6 concentration, using the affinity parameters from Figure 3, shown as a red dashed line and plotted on the right y-axis.

The relative *k*_+_ in the presence of varying JB6 concentrations are obtained from the initial slopes of the aggregation reactions in Figure 2d, and divided by k_+_ of Aβ42 alone. This is used as a measure of the inhibition potency.

The JB6 saturation level of the pre-formed fibrils (seeds) at time zero of the aggregation process is calculated with Equation 1 and 2 using the affinity parameters determined in Figure 3a-b (K_D_ = 1 nM and a stoichiometry corresponding to approximately 1:7 JB6 to Aβ42 monomers in fibrils). At time zero there are 250 nM fibrillar and 4.75 µM monomeric Aβ42 and the maximal amount of monomeric JB6 that can bind to fibrils can thus be calculated as 250/7 nM = 36 nM, which is defined as a saturation level of 1.0 in Figure 4.

The calculated saturation level of the fibrils at each JB6 concentration displays a trend very similar to the relative *k*_+_, indicating that the binding of JB6 is directly related to the inhibition level. Moreover, this finding implies that the affinity for the fibril end is similar to that of the general fibril surface.

At JB6 concentrations above approximately 100 nM, not all JB6 is monomeric. However, the monomer concentration is around 100 nM, which is sufficient to provide near complete fibril saturation. The remaining JB6 may bind to newly formed aggregates as monomers become available over time through ongoing micelle dissociation [38], further prolonging the apparent lag phase. This explains why t_1_*_/_*_2_ increases linearly with JB6 concentration, as seen both in Figure 2c and 1d.

Figure 5 illustrates how crowded the fibril surface would be at the saturation point of 1 JB6 monomer bound per 7 Aβ42 monomers in the fibril. For this illustration we have used a fibril structure determined using a combination of solid state NMR (ssNMR) and small angle X-ray scattering (SAXS), which have four monomers in the cross-sectional plane [44]. The JB6 molecules are added onto the fibril in the form of identical copies of a monomeric JB6 structure predicted using AlphaFold3 [49, 50].

**Figure 5:**
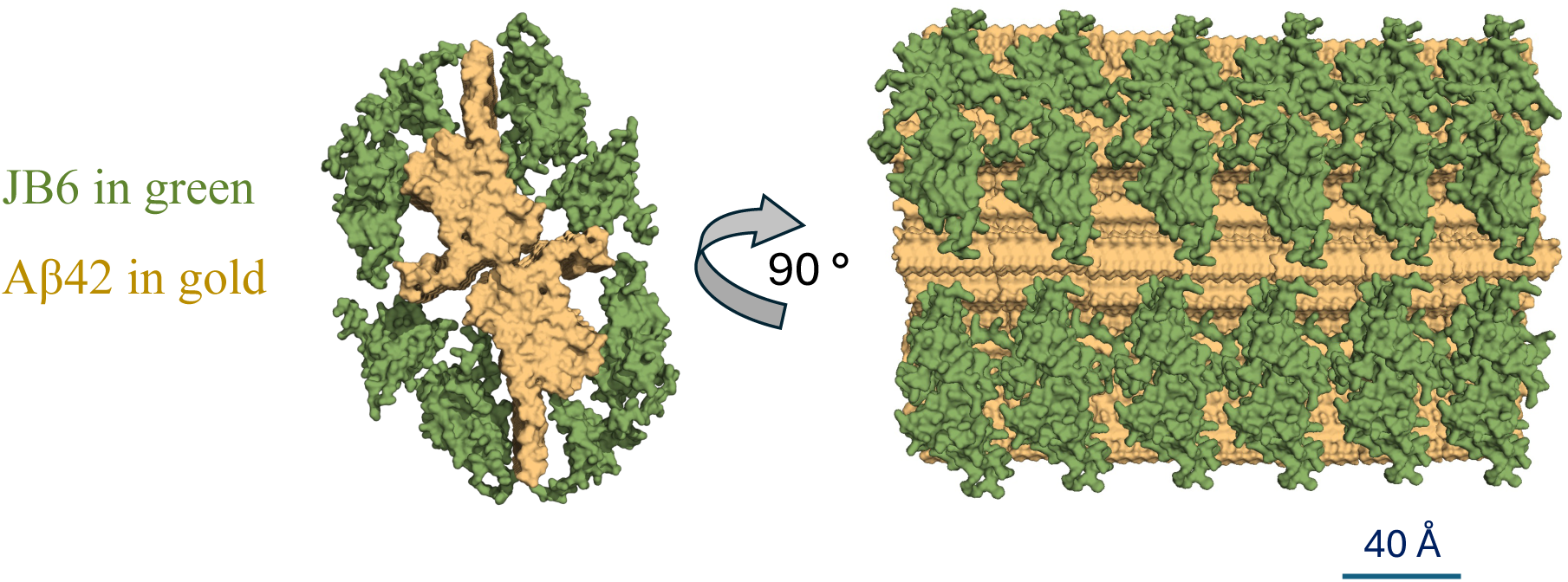
Visualization of the molar ratio obtained from the affinity measurements. 1 JB6 molecule to 7 Aβ42 molecules (in fibril state). Aβ42 is shown in gold and JB6 in green. The same complex is shown from two perspectives. No structural information should be inferred from this illustration, except for the Aβ42 fibril structure itself, which is solved using a combination of ssNMR and SAXS [44]. This fibril structure consist of four Aβ42 molecules in the cross-sectional plane, two in each filament. The JB6 structure is predicted using AlphaFold3 and simply placed next to the fibril, for visualization purpose. The figure was generated in PyMol using surface representation.

Note that the illustration is only intended to visualize the occupied volume at a 1:7 molar stoichiometry and no structural information should be inferred from the arrangement shown. However, the molecular volumes themselves are independent of the detailed structure of the bound molecules, and the illustration can therefore help in reasoning about the JB6 binding to Aβ42 fibrils. For instance, it appears unlikely that JB6 only binds to isolated defects along the fibril, as has been inferred for Brichos domains with significantly lower stoichiometry [51]. Rather, the relatively high stoichiometry of JB6 binding suggests that the fibril surface is essentially coated by JB6. A similar coating has been found in the case of DNAJB1 binding to α-synuclein fibrils [52], with an arrangement and spacing that resembles the illustration in Figure 5. Similar coating of Aβ42 fibrils has been observed for the antibody aducanumab [53].

We note that the 95 % confidence interval of the *K_D_* determination of JB6-fibril binding ranges between 0 and 2.6 nM. This is because the JB6 concentration is about 20 nM, and the experiments are therefore less sensitive to higher affinities (lower *K_D_*). Thus, we cannot exclude that there may be additional binding sites with higher affinities and very low concentration of binding sites, for example fibril defects which are found to be crucial for secondary nucleation [51].

### Probing JB6 interactions during the aggregation using diffusion-based measurements

To further understand what determines the length of the JB6-dependent lag phase observed in Figure 1 and 2, we withdrew samples at multiple time-points during the aggregation process and analyzed the JB6 aggregation state (with workflow as illustrated in Figure 6a). A non-labeled aggregating system (Figure 6b) with 5 % sonicated seeds and a total Aβ42 concentration of 5 µM and 100 nM JB6 was probed using CD spectroscopy (Δ*ɛ*_218nm_). After centrifugation of samples withdrawn during the reaction, the concentrations of JB6 and Aβ42 were measured using HPLC. Interestingly, both JB6 and Aβ42 are kept from forming large aggregates during the entire lag phase, but at the end of the lag phase both proteins disappear from the supernatant, mirroring the amyloid formation kinetics. Non-normalized data are shown in Figure S4b, together with an analysis of the concentrations in relation to the affinity parameters obtained in Figure 3, as discussed in supplementary note 1.

**Figure 6:**
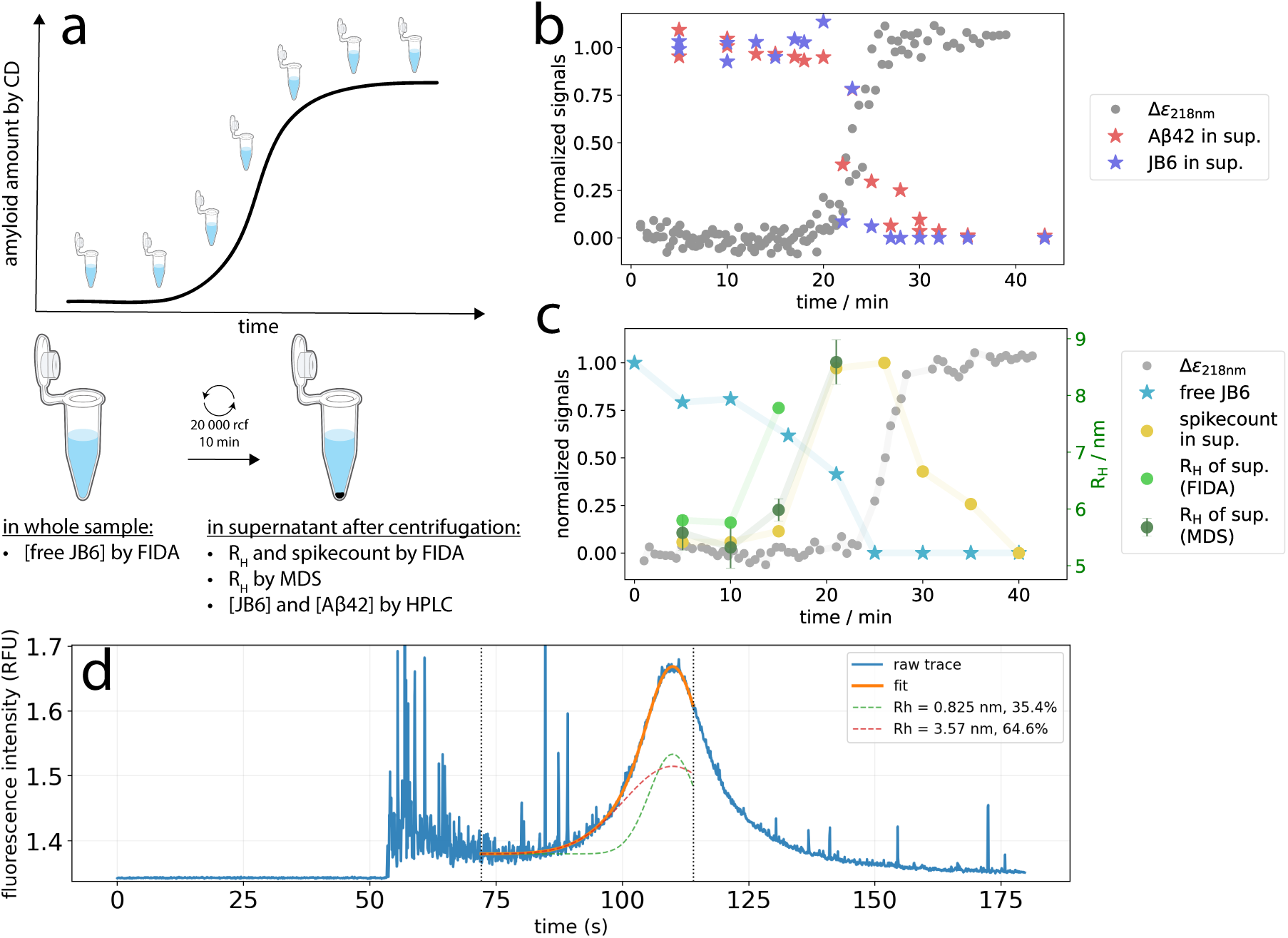
Probing supernatant concentrations and complex sizes during amyloid formation to better understand the inhibition mechanism and aggregation kinetics. **a:** Experimental workflow. Aggregation was monitored by CD spectroscopy and samples were withdrawn during the reaction and analyzed by FIDA to quantify free (fluorophore-labeled) JB6. A centrifugation step removed large aggregates and the supernatant was analyzed by MDS and FIDA for size determination of JB6 and JB6-complexes. HPLC was used to quantify Aβ42 and JB6 concentrations in the supernatant. The starting point was 5 % sonicated seeds, 5 µM total Aβ42, and 100 nM JB6. b: Non-labeled proteins to obtain concentrations of Aβ42 and JB6 in the supernatant during the aggregation. The normalized Δɛ218nm signal reports on β-sheet content and thus amyloid formation. c: As in (b), but with fluorophore-labeled JB6 to determine JB6 size and complex formation. Δɛ218nm, free JB6 concentration, and spike count are shown on a normalized y-axis. Spike count reports on larger particles not resolved by Taylor dispersion under these conditions (400 mbar). The hydrodynamic radii (obtained using MDS and FIDA) are shown on the right y-axis. d: Raw FIDA-data showcasing spikes and fit with two species. The ∼1 nm specie is free fluorophore and the 3-4 nm specie is free JB6 [38]. This Taylor dispersion profile correspond to 10 minutes into the aggregation process in panel (c), before centrifugation.

So, why does the lag phase suddenly end when both proteins are still present in the supernatant? What differs at this point compared to the start of the reaction? To address this, we used fluorophore-labeled JB6 and quantified its diffusivity to estimate the size of JB6 and its complexes (Figure 6c). Free JB6 was quantified using flow-induced dispersion analysis (FIDA, [54, 55]). See Figure 6d and S5–S6 for examples of FIDA data and analyses on samples during the aggregation process. The monomeric JB6 population decreases during the lag phase and approaches zero at its end (Figure 6c). In parallel, the apparent size of fluorescent species in the supernatant increases, indicating formation of co-oligomers. The size of these species exceed the quantification limits of MDS (*∼*20 nm) and FIDA (*∼*13 nm at 400 mbar). Instead, FIDA detects them as spikes, reported as the semi-quantitative measure “spike count” as showcased in Figure 6d as the raw data Taylor dispersion profile of the sample at 10 minutes into the aggregation process. Such complexes displays a peak in abundance near the half-time of aggregation, as expected of amyloid oligomers [33] and consistent with previous observations of JB6 binding to Aβ oligomers [16, 19]. The complexes remain small enough to avoid pelleting during centrifugation, explaining the constant JB6 concentration in the supernatant (Figure 6b).

To summarize the findings in Figure 6, JB6 renders the seeds practically incompetent of elongation and secondary nucleation, resulting in a lag phase. During this period, the nucleation of new fibrils occurs in a dynamic process with a majority of oligomers failing to convert and instead dissociating [33]. JB6 binds to these transient species, reducing their rate of conversion/nucleation and thus limiting the following rapid growth. Since JB6 is absent in the supernatant after 40 min, the JB6–Aβ42 complexes either grow, associate with fibrils, or dissociate during the growth phase and early plateau. Once close to all JB6 is occupied in complexes with Aβ42, the bulk amyloid formation cannot be delayed further and rapid aggregation ensues, as has been suggested before [11], and similar to observations in inhibition of tau aggregation [14]. This is supported by the similar aggregation kinetics of the second half of the aggregation process being independent of the JB6 concentration, as shown in Figure S1 and consistent with findings in Figure 2e; the first part of the reaction is strongly JB6 dependent in both slope and length, but the second part is independent of JB6 concentration (for JB6 concentrations < cmc).

Reducing the Aβ42 concentration by half slows the process (Figure S7) as expected [31] but yields similar behavior with an extended lag time and observations of JB6-Aβ42 complex-formation with maximal prevalence around t_1_*_/_*_2_. However, the complexes are smaller (*∼*10 nm radius) and remain within the Taylor-dispersed regime.

### Conclusions

In conclusion, JB6 inhibits Aβ42 elongation if added to pure pre-formed Aβ42 fibrils, complementing its established suppression of primary and secondary nucleation. Together, these effects enable JB6 to render fibrils inactive for further growth while preventing the formation of new nuclei, indicating that JB6 is a complete amyloid suppressor with strong potential for therapeutic intervention. The measured affinity of JB6 for Aβ42 fibrils directly links its binding to the inhibition of amyloid growth, providing a quantitative framework for chaperone-mediated suppression. This work may be extended to similarly describe inhibition of primary and secondary nucleation and represents a step toward modeling amyloid inhibition based on binding properties. Finally, JB6 binds to small Aβ42 intermediates (amyloid oligomers) or forms co-oligomers, and when close to all chaperone is bound, aggregation ensues with same kinetics as without JB6.

## Methods

All chemicals were of analytical grade. All buffers were filtered through a wwPTFE-filter with 0.22 µm pore size and degassed.

### JB6 and Aβ42 expression and purification

Both proteins were expressed in the *E. coli* strain BL21 DE3 pLysS star, and purified in a tag-free manner using the protein’s biophysical properties, as described in detail in [31, 56, 57].

JB6, with the sequence MVDYYEVLGVQRHASPEDIKKAYRKLALKWHPDKNPENKEEAERKFKQVAEAYEVLSDAKKRDIYDKYGKEGLNGGGGGGSHFDSPFEFGFTFRNPDDVFREFFGGRDPFSFDFFEDPFEDFFGNRRGPRGSRSRGTGSFFSAFSGFPSFGSGFSSFDTGFTSFGSLGHGGLTSFSSTSFGGSGMGNFKSISTSTKMVNGRKITTKRIVENGQERVEVEEDGQLKSLTINGKEQLLRLDNK, was flash-frozen after purification using pre-cooled metal blocks and tubes (-80 °C), then stored at -20 °C until thawing in 37 °C heatblock prior to use.

Aβ42, with the sequence MDAEFRHDSGYEVHHQKLVFFAEDVGSNKGAIIGLMVGGVV IA, was lyophilized after purification, stored at -20 °C until usage, dissolved in 6 M guanidine (20 mM NaP buffer, pH 8.0) and run on a superdex 75 size exclusion chromatography (SEC) column in the experimental buffer. The isolated monomers were kept on ice until usage, which was within a few h.

NCysJB6, with the sequence MCVDYYEVLGVQRHASPEDIKKAYRKLALKWHPDKNP ENKEEAERKFKQVAEAYEVLSDAKKRDIYDKYGKEGLNGGGGGGSHFDSPFEFGFTFRNPDDVFREFFGGRDPFSFDFFEDPFEDFFGNRRGPRGSRSRGTGSFFSAFSGFPSFGSGFSSFDTGFTSFGSLGHGGLTSFSSTSFGGSGMGNFKSISTSTKMVNGRKITTKRIVENGQERVEVEEDGQLKSLTINGKEQLLRLDNK, was expressed and purified in the same manner as wt JB6 except that 1 mM DTT was included in all buffers. The protein was flash-frozen after purification using pre-cooled metal blocks and tubes (-80 °C), then stored at -20 °C until thawing in 37 °C heatblock prior to use. Before labeling, DTT was removed using SEC in 20 mM NaP, pH 8.0, after which 2 molar equivalents of CF647 were added to the protein from a concentrated stock. The solution was incubated for 5 h at room temperature, lyophilized, dissolved in 1 ml of 6 M GuHCl and excess dye was removed by SEC. An absorbance spectrum was recorded for the collected CF647-JB6 using a Labbot instrument and a 10 mm cuvette. The concentrations of protein and dye were calculated from the absorbance at 280 and 647 nm, 0.039 and 0.335, respectively, using extinction coefficients of 240,000 l mol*^−^*^1^ cm*^−^*^1^ at 647 nm and 7,200 l mol*^−^*^1^ cm*^−^*^1^ at 280 nm for the dye, and 14400 l mol*^−^*^1^ cm*^−^*^1^ at 280 nm for the protein, resulting in 1.4 µM CF647 and 2 µM protein and an estimated degree of labeling of 0.7 dye per protein.

### Affinity measurements using HPLC

Aβ42 fibrils were prepared from 10 µM monomers, with a magnetic stirring bar (500 rpm) at 37 °C for 3 h. The buffer used was 20 mM NaP, 0.2 mM EDTA, pH 8.0. JB6 was pre-equilibrated at 1.5 times the final concentration (in 5 ml Eppendorf protein low-binding tubes, in an initial volume of 5 ml), at 37 °C for 3 h, before being mixed with pre-formed fibrils at a sample volume of 200 µl in 1.5 ml Eppendorf protein low-binding tubes. The samples were let to incubate overnight (17 h) at 37 °C, followed by centrifugation in the same tubes at 20 000 g for 30 min. The top 100 µl of supernatant was analyzed with HPLC (Shimadzu Nexera X3) to measure the JB6 concentration.

In one series of samples, the JB6 concentration was kept constant and the fibril concentration was varied (with concentration representing the monomer equivalent). All samples had the same volume of the same JB6 solution, providing the same JB6 concentration (calculated to 40 nM). Due to surface activity of JB6 [14, 48], the actual value of this concentration is measured (quantified as a fitted parameter), rather than calculated based on the stock concentration and dilution factor. In another series of samples, the JB6 concentration was varied and the fibril concentration kept constant (100 nM). Since JB6 adsorption to the tube surfaces is concentration dependent [14, 48], the actual JB6 concentration in each sample was measured in an exact duplicate sample but with added buffer instead of fibrils. The difference in JB6 concentration in this sample and the sample with added fibrils is the fibril-bound amount of JB6.

A global fit to both data sets was performed with fitted global parameters being K_D_, and a stoichiometry factor at saturation, *s*, and a local parameter being the JB6 concentration in the data set with constant JB6 concentration.

The HPLC analysis method and settings: 40 µl of the sample was loaded on a 60 °C C18 reversed phase column (BIOshell A160 Peptide CN column 66966-U, Sigma-Aldrich). Elution was done with a 10 minutes linear gradient of 5 to 95 % acetonitrile in an aqueous mobile phase, with 0.1 % TFA, at 0.5 ml/min. Absorbance at 280 nm was measured and used to quantify the protein concentration, using the extinction coefficient of 14440 cm*^−^*^1^M*^−^*^1^ for JB6 and 1490 cm*^−^*^1^M*^−^*^1^ for Aβ42. The plate used for the HPLC auto-sampler was a PEGylated low-binding 96-well plate, half area (3686, Corning). It was found to be important not to keep the plate in the auto-sampler chamber at as low temperature as 10 °C, since loss of JB6 (likely due to surface adsorption) was found to be much more pronounced compared to at 25 or 37 °C, as investigated in supplementary information, S8. For the present work, the auto-sampler chamber was kept at 25 °C.

### Aggregation kinetics using ThT fluorescence

Freshly isolated Aβ42 monomers were mixed with ThT and kept on ice until usage. The buffer was 5 mM NaP, pH 8.0, 0.02 mM EDTA, 40 mM NaF (low absorbance in CD spectroscopy, but similar ionic strength as 20 mM NaP). Seeds were prepared in the same buffer, with the same concentration ThT used in the aggregation kinetics (5 µM), with 10 µM Aβ42, and magnetic stirring bar (500 rpm) at 37 °C for 1 h, followed by sonication with a tip sonicator for 5 minutes (1 sec on, 1 sec off), in an ice slush to keep the temperature low.

Since JB6 needs to dissociate into monomers to be fully active as inhibitor [18], and the dissociation occurs with a halftime of about 20 minutes at 37 °C [38], JB6 was pre-diluted to 1.5 times the final concentration directly in the plate (PEGylated low-binding 96-well plate, half area (3881, Corning)). The plate was incubated with lid for 2 h at 37 °C before addition of seeds, and two minutes later monomeric Aβ42 was added with multichannel pipette. The total volume in each well was 100 µl, with three wells of each condition, with plastic cover to prevent evaporation. The aggregation was probed with continuous reading (excitation at 380 nM and emission recorded at 480 nM), cycle time = 29 s, with a platereader (FLUOstar Omega, BMG LABTECH) at 37 °C.

### Aggregation kinetics using CD spectroscopy

CD spectroscopy (with a J-815, JASCO) was performed in a quartz cuvette with 4 mm path-length, at 37 °C (adjusted with a Peltier temperature controller). For the spectra, the measured range was 260-195 nm, at a scanning speed of 50 nm/min, 8 s D.I.T., 1 nm bandwidth, and an average of 3 accumulations. When probing the ellipticity at 218 nm, 5 µM Aβ42, with 5 % seeds (formed from a 12 µM solution in the CD spectrometer, with the same solution conditions), and varying JB6 concentrations, were used. To keep the sample well dispersed during the whole experiment, a magnetic stir bar was used at a slow but continuous rotation. The experiments were run in sequence, with a thorough wash of the cuvette in between (soak in Hellmanex III for 10 minutes and rinse with MilliQ water 10 times), with Aβ42 on ice until usage. The buffer was 5 mM NaP, pH 8.0, 0.2 mM EDTA, 40 mM NaF (which has a low absorbance in CD spectroscopy at around 200 nm). The JB6 contribution to the CD signal at 218 nm was measured at each JB6 concentration (shown in Figure S9), providing a baseline to be subtracted in the data of Figure 2 and Figure S3. Also in this experiment JB6 was pre-equilibrated to 1.5 times the final concentration and mixed with seeds for two minutes before addition of the monomeric Aβ42.

### Concentration and size determinations during aggregation

#### CD spectrometry

The same conditions and CD spectrometry methodology were used as described in “Aggregation kinetics using CD spectroscopy”. 100 nM JB6 was used (pre-equilibrated at 150 nM at 37 °C for 3 h) together with 0.25 µM Aβ42 seeds and 4.75 µM monomers.

#### HPLC to quantify protein concentrations in supernatant

In one set of experiments, non-labeled JB6 was used to measure the JB6 and Aβ42 concentrations in the supernatant after centrifugation at 20 000 rcf for 10 min. 100 µl of the ongoing reaction in the CD spectrometer was transferred to an Eppendorf protein low-binding tube for centrifugation. 50 µl of the surface layer was transferred to a PEGylated 96-well plate (Corning, 3686), kept at 25 °C to minimize surface adsorption, and analyzed using HPLC (Shimadzu Nexera X3) with the same settings as described in “Aggregation kinetics using ThT fluorescence”.

#### Particle sizing using FIDA and MDS

In a second set of experiments, fluorescently labeled JB6 (CF647-JB6) was used to measure the diffusivity, to size JB6 and its complexes, in addition to the CD spectroscopy and HPLC quantification. After pre-equilibrating CF647-JB6 at 37 °C overnight at 150 nM, CD spectroscopy and HPLC were performed as described above. Prior to centrifugation, 9 µl of the reaction solution was mixed with 1 µl of 0.3 % tween and analyzed immediately with FIDA (FIDA Neo, Fidabio, 640 nm detector), using a permanently hydrophilic-coated capillary at 25 °C. The low sample volume was enabled by using pressure vial inserts from Fidabio. The measurements were run with 400 mbar for mobilization and measurements, with the sample buffer as analyte (including 0.03 % tween) and sample as indicator. After centrifugation, the supernatant was analyzed again with FIDA, with a delay of 60 minutes due to the ongoing measurements during the aggregation process. However, 10-20 minutes after centrifugation, the supernatant was analyzed with MDS, using a Fluidity One M (Fluidic Sciences, Cambridge, UK) in size setting 3 and viscosity setting 1, in technical triplicates. Additionally, HPLC was used to quantify the protein concentrations in the supernatant as described for non-labeled JB6.

The hydrodynamic radius was extracted from the FIDA data by fitting the peak-excluded raw data to Equation 3, which describes Taylor dispersion:

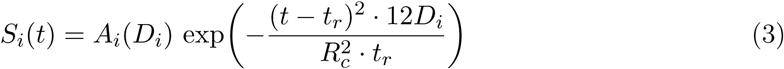

Where *S_i_* is the fluorescence signal of specie *i*, *t* the time of flow in the capillary, *A_i_* the component amplitude, *D_i_* the diffusion coefficient, relating to the hydrodynamic radius, *R_H_* via the Stokes-Einstein relation: 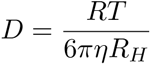, *t_r_* the time at the peak maximum, and *R_c_* the capillary inner radius (37.5 µm). Examples of raw data and fits are shown in Figure S5-S6.

## Acknowledgments and funding sources

Swedish Research Council grant 2015-00143 (SL)

European Research Council grant 101097824 (SL)

Knut and Alice Wallenberg Foundation grant 2022-0059 (SL, UO).

