## Supplemental Information for "DNAJB6b — a complete amyloid inhibitor"

**A. Carlsson, U. Olsson and S. Linse**

### Supplemental note 1

In the equation describing growth via elongation: $(f(t) = 1 - 0.95e^{-kt})$, $k$ relates to the elongation rate constant, $k_+$, as $k = k_+2[P]$, where $[P]$ is the number concentration of fibrils, and $2[P]$ is the number concentration of fibril ends, which is to a good approximation constant in the seeded reactions we examine in this work. Thus, the exact value of $k_+$ is not of high importance here, since the $k$-value is experimentally obtained and requires less assumptions compared to $k_+$. However, as a quality control we can use some reasonable assumptions to calculate $k_+$ and compare with what has previously been reported.

$k$ was in Figure 1a found to be 0.00182±0.00009 $s^{-1}$. $[P]$ was estimated to 0.6 nM, based on $0.05 \times 5$ μM = 0.25 μM sonicated seeds with an estimated fibril length of 50 nm [41, 42, 43], 4 monomers per plane [44], with 4.7 Å spacing between planes. This gives $k_+ = 1.5 \times 10^6$ $M^{-1}$ $s^{-1}$, which is only a factor of two lower than previously determined [34], which is considered within the expected error margin.

### Supplemental note 2

Figure 2c presents an example of how much JB6 is bound to a container's surfaces, and how much is bound to fibrils. If the surface adsorption would not be measured separately, one could be led to conclude that all loss of JB6 is to fibrils, and thus overestimate the bound amount greatly. This particular example comes from an experiment of 100 nM JB6, and 250 nM Aβ42 fibrils, which is the initial state before 4.75 μM Aβ42 monomers was added and aggregation kinetics was probed with CD spectroscopy, with data shown in Figure 6b.

Both samples (JB6 alone and JB6 + fibrils) encountered the same surfaces and steps: mixing in the cuvette used for the aggregation reaction in the CD spectrometer (at 37 °C), transferring 100 μl to a Eppendorf protein lo-bind tube and centrifuging for 10 min at 20 000 g, transferring 50 μl of the top layer to a PEGylated non-binding 96-well plate (Corning, 3686), kept at 25 °C, and analyzing the concentration in HPLC reversed phase. The difference in JB6 concentration between the two samples is the fibril-bound amount of JB6.

This difference is 43±5.3 nM −13±2.5 nM = 30±8.3 nM. Using the affinity parameters determined in Figure 3 and Equation 1 and 2, the amount of bound JB6 should be 37 nM, which is considered well within the experimental uncertainty.

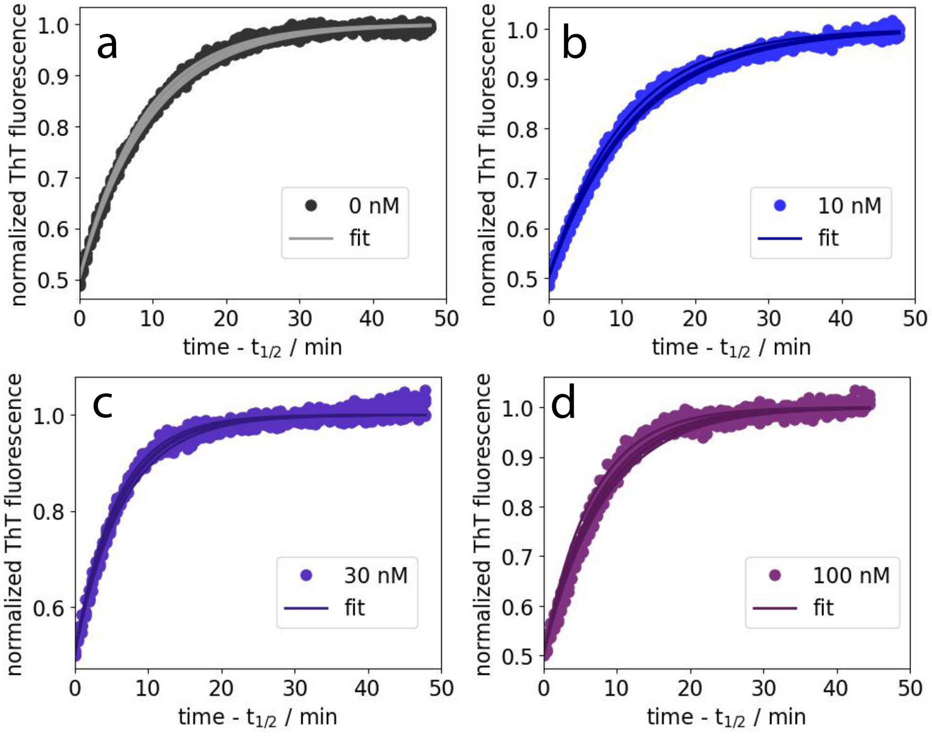


Figure S1: The final 50 % of the kinetic traces from Figure 1, fitted to a model with only elongation (as described in the main text, in conjugation to Figure 1c). **a-d**: 0, 10, 30, and 100 nM JB6. This analysis is applicable for the JB6 concentrations below its cmc, since above this concentrations the micelles can dissociate and thus continue to affect the aggregation also in the later stage of the aggregation process.

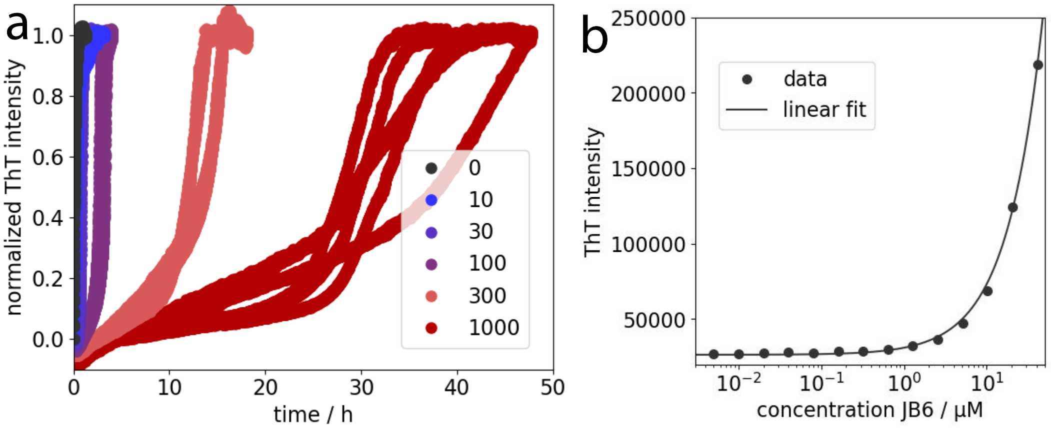


Figure S2: Supplemental information to Figure 1. **a**: Same data as in Figure 1b but during 50 h to show that Aβ42 aggregates even in the presence of 300 and 1000 nM JB6. **b**: ThT fluorescence as a function of JB6 concentration in 20 mM NaP buffer, 0.2 mM EDTA, pH 8.0, with a linear regression showing a fairly linear correlation. Note that the x-axis is in logarithmic scale. The ThT concentration was kept constant at 5 µM while JB6 was varied from 4 nM to 40 µM.

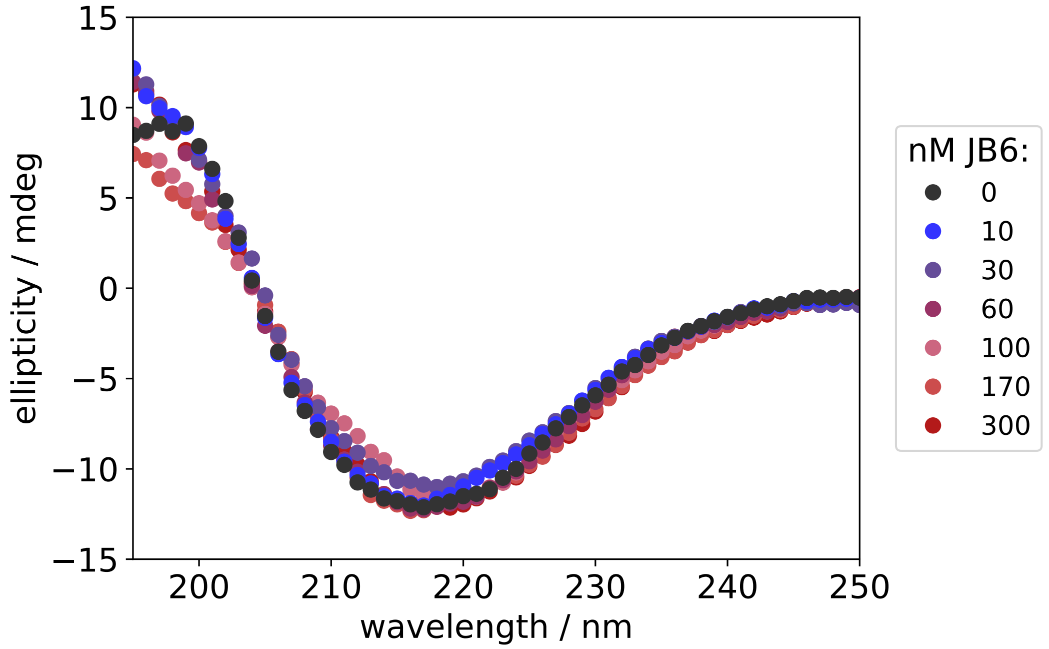


Figure S3: CD spectra on end-states in the aggregation reactions in Figure 2b.

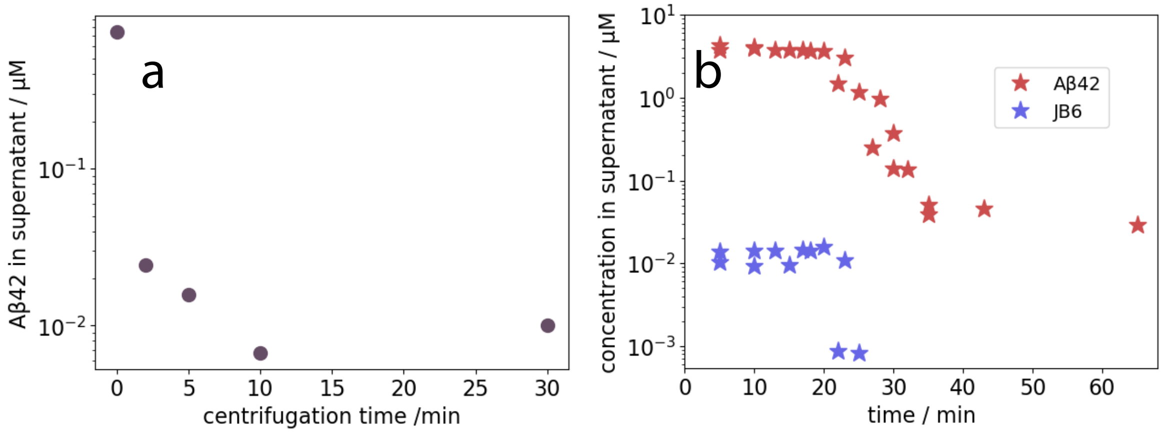


Figure S4: Concentration measurements supporting Figure 6. **a**: Aβ42 concentration in the supernatant as a function of centrifugation time with 20 000 g. 10 min is enough to sediment most of the fibrils. **b**: Concentrations of Aβ42 and JB6 in absolute scale (logarithmic) during the aggregation process. The Aβ42 concentration plateaus at about 30 nM whereas the JB6 concentration is lower than the limit of detection (less than 1 nM) after the aggregation and is therefore not plotted.

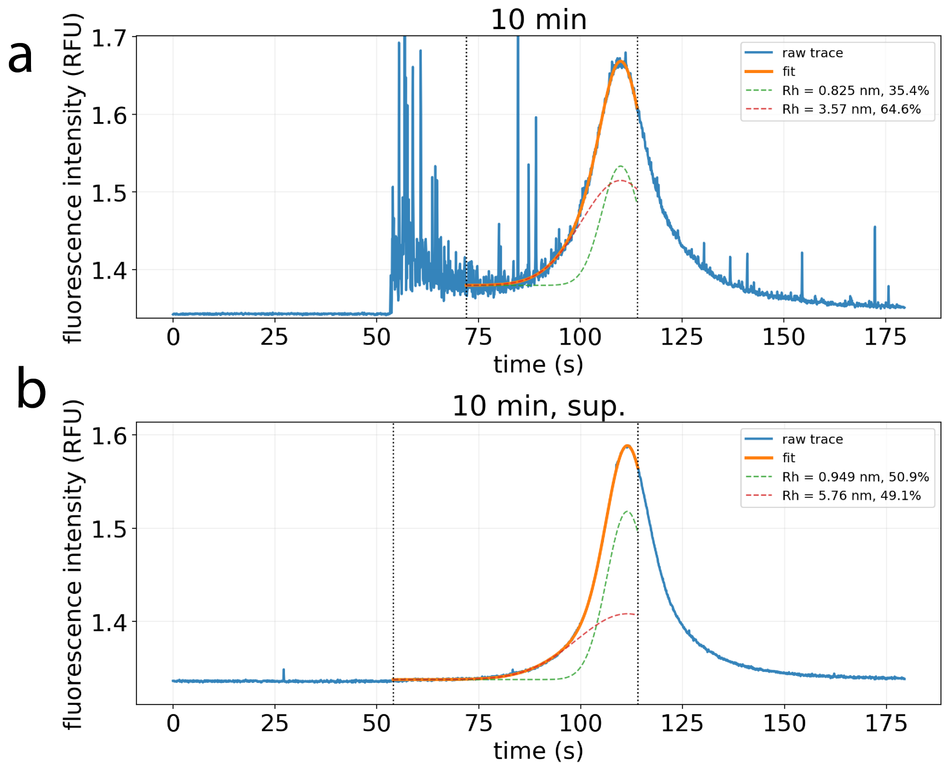


Figure S5: Raw FIDA data of the sample 10 minutes into the reaction of Figure 6c, before (**a**) and after (**b**) centrifugation. The many spikes in (a) are expected since there are fibrils present in the sample. After excluding the spikes, the data in the window between 72-114 s for (a) and 54-114 s for (b) were used to fit Equation 3, with two species superimposed. As can be seen one component is around 0.8-1 nm in radius, which is explained as some free CF647 dye in the sample. But this is clearly distinct from the 3-4 nm radius of monomeric JB6 [38] and therefore the amount of free JB6 can be extracted in this way.

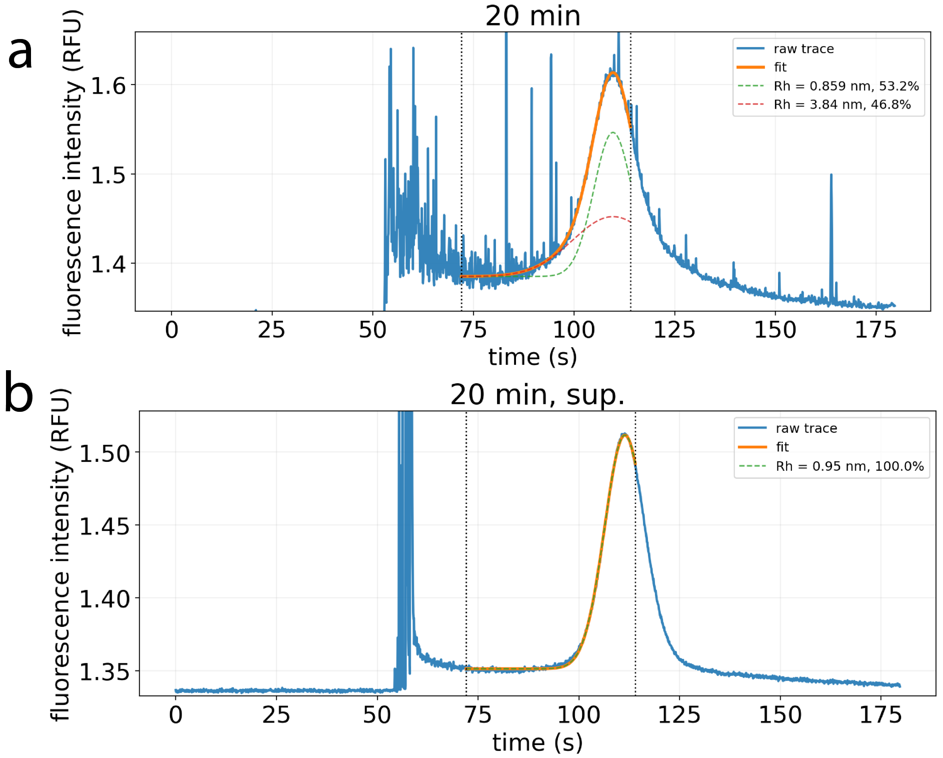


Figure S6: Raw FIDA data of the sample 20 minutes into the reaction of Figure 6c, before (**a**) and after (**b**) centrifugation. In this case, the complexes in the supernatant are so large they end up as spikes instead of in the peak, which is why the measure "spikecount" was used to semi-quantitatively probe the amount of complexes in the supernatant as shown in Figure 6c.

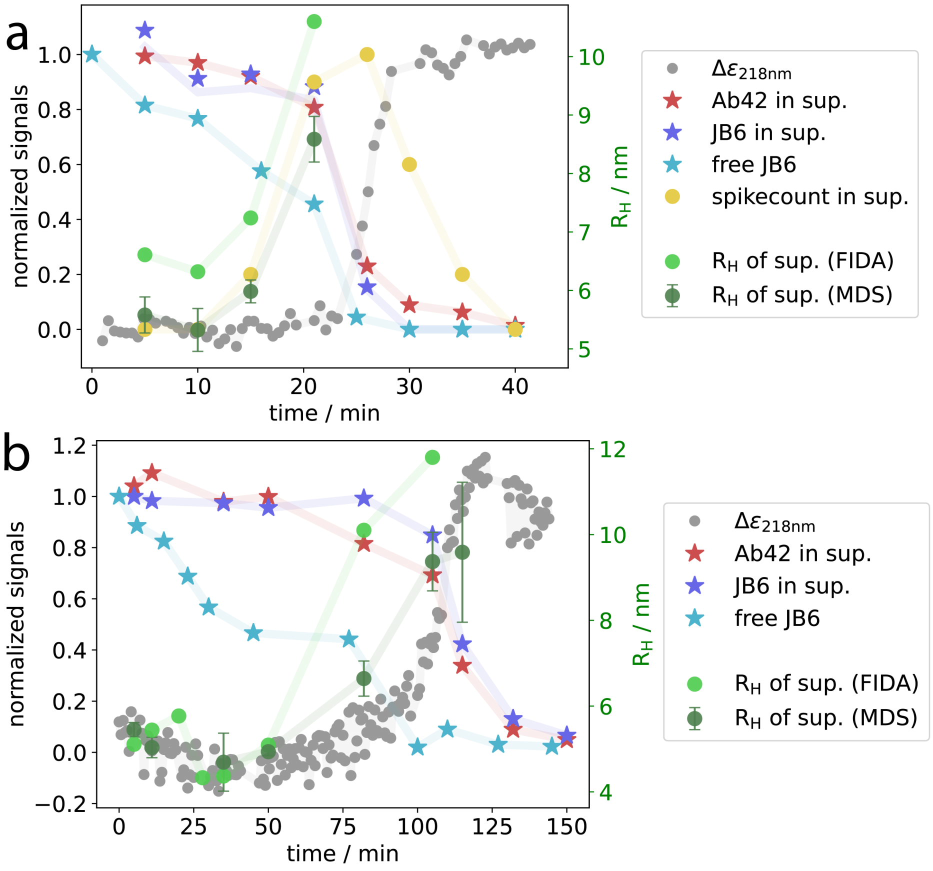


Figure S7: Probing protein concentrations in the supernatant, free JB6 in the sample, and size of JB6-complexes during aggregation of 5 and 2.5 µM total Aβ42 concentration, with 0.125 µM respectively 0.25 µM seeds. **a**: Same data as in Figure 6c but including also concentrations of both Aβ42 and JB6 in the supernatant, measured with HPLC. **b**: Same methodology but with half the Aβ42 concentration to obtain slower kinetics, enabling more data points, and the longer time in between data points made it possible to run FIDA on the supernatant shortly after the centrifugation. Therefore, the complexes in the supernatant are better sized and end up in the Taylor dispersed region instead of as spikes.

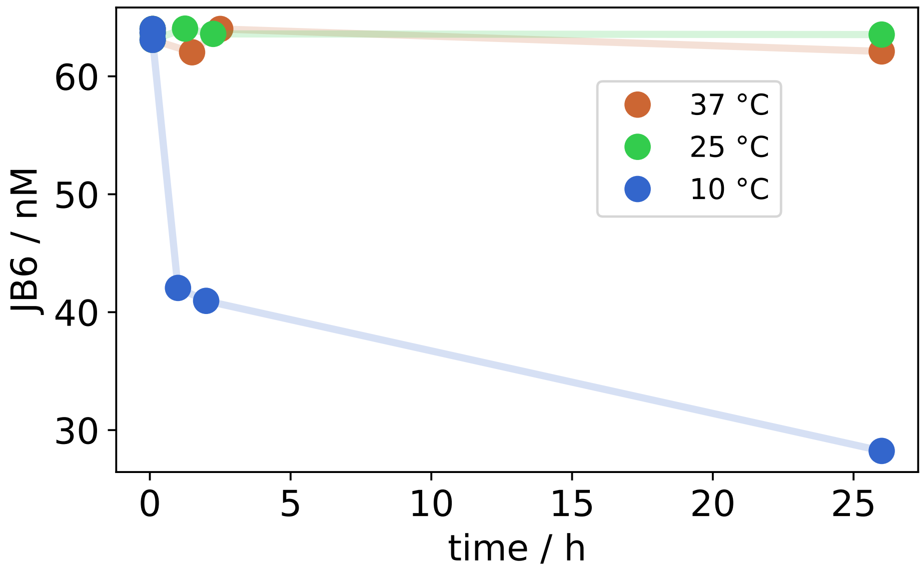


Figure S8: JB6 losses in a PEGylated low-binding 96-well plate, half area (3686, Corning), at three temperatures during the incubation time. At 25 and 37 °C, the loss of protein at surfaces is smaller than the variance of the measurement. At 10 °C, the loss of protein is dramatic and will provide a large problem when the plate is incubated in the auto-sampler of the HPLC. Thus, a higher temperature should be used for this purpose (at least in the case of JB6).

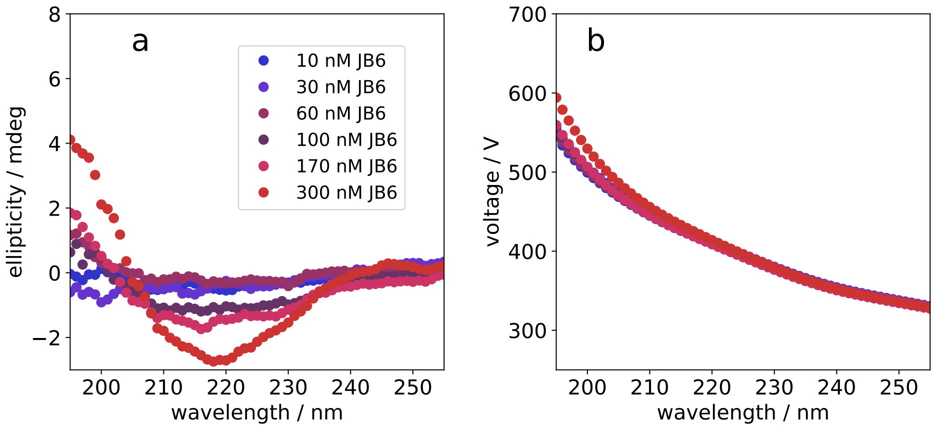


Figure S9: JB6 at various concentrations measured with CD spectroscopy. **a**: Baseline subtracted ellipticity. **b**: The automatically generated voltage used to amplify the signal, which preferably should be below 600 V to ensure good quality data.
